# Tau forms dynamic hot spots that are resistant to microtubule perturbations and cholesterol depletion

**DOI:** 10.1101/2022.02.04.479198

**Authors:** Pranesh Padmanabhan, Andrew Kneynsberg, Esteban Cruz Gonzalez, Rumelo Amor, Jean-Baptiste Sibarita, Jürgen Götz

## Abstract

Accumulation of Tau aggregates is a pathological hallmark of Alzheimer’s disease, but little is known about where these aggregates assemble and how stable these assemblies are. Using quantitative, single-molecule imaging, we show that Tau exhibits spatial and kinetic heterogeneity near the plasma membrane, resulting in the formation of nanometer-sized hot spots smaller than the diffraction limit. The hot spots lasted tens of seconds, much longer than the short dwell time (~40 ms) of Tau on microtubules. Pharmacological and biochemical disruption of Tau/microtubule interactions did not affect hot spot formation, suggesting that these are different from the reported Tau condensation on microtubules. Although cholesterol removal has been shown to reduce Tau pathology, its depletion did not affect Tau hot spot dynamics. Our study identifies an intrinsic dynamic property of Tau near the plasma membrane that may facilitate the formation of assembly sites for Tau to assume its physiological and pathological functions.

## INTRODUCTION

Tau is conventionally thought of as a protein that primarily associates with microtubules, thereby regulating their stability^1–3^. However, as an intrinsically disordered protein, Tau has remarkable structural flexibility, and it also undergoes various post-translational modifications, allowing it to interact with numerous binding partners and execute multiple cellular functions^1–3^, including axonal transport^4–6^, neuronal maturation^7,8^, and protein translation^9,10^. Interest in the behavior of Tau at the plasma membrane was spurred firstly by the realization that, in an Alzheimer’s disease (AD) context, toxic Tau aggregates have been reported to cross the plasma membrane for release into the extracellular milieu, thereby contributing to the spread of Tau across interconnected brain regions, a process believed to underlie the pathology in AD and other tauopathies^1,11^. Secondly, in an ill-defined physiological context, Tau lacking a classical signal peptide undergoes unconventional secretion^12,13^.

The association of Tau with the plasma membrane can occur directly through electrostatic interactions with anionic lipids^14,15^ and indirectly through interactions with membrane-associated proteins, such as the Src kinase Fyn^16^, the motor protein dynactin^17^, and the phospholipid-binding protein annexin A2^18^. Moreover, Tau interacts with microtubules and actin filaments, anchoring them to the plasma membrane^19^. The question arises whether Tau forms aggregates near the plasma membrane, and if so, how stable these assemblies are. So far, studies using artificial lipid membrane studies have suggested that membrane binding can cause conformational changes to Tau^20^, initiate Tau aggregation and fibrilization^21,22^, and mediate Tau-induced membrane permeabilization and toxicity^23,24^. However, despite the critical importance of Tau-membrane interactions^12,13,16,25,26^, quantifying how Tau behaves and organizes at the plasma membrane in a live cellular environment has previously proven challenging because of the lack of sensitive tools, thereby limiting our understanding of how this protein executes its physiological and pathological functions in this compartment^27^.

Here, using single-molecule-based super-resolution microscopy, we investigated both the dynamics and nanoscale organization of Tau near the plasma membrane of live cells. We found that Tau exhibited immobile, confined and free diffusive states and formed nanometer-sized hot spots. Moreover, Tau confinement and immobilization were due to its trapping on microtubules and in the hot spots. We found that while inhibiting Tau/microtubule interactions released Tau from microtubules and increased Tau’s mobility, the heterogeneous mobility pattern persisted and surprisingly, the dynamics of Tau hot spot was unaltered. Moreover, these hot spots were also resistant to depletion of cholesterol, which is known to reduce Tau pathology in transgenic mice^28^. Here, we suggest a model whereby the dynamic partitioning of Tau into different kinetic pools equips Tau with the dynamic properties to form assemblies near the plasma membrane as a critical step in pathological dysregulation.

## RESULTS

### Single-molecule tracking of Tau near the plasma membrane

To visualize and track individual Tau molecules near the inner leaflet of the plasma membrane, we transiently expressed the most prevalent human 0N4R isoform of Tau (383 amino acids) fused carboxy-terminally to photoactivatable fluorescent protein mEos3.2 (hereafter abbreviated to Tau^WT^-mEos3.2) in murine N2a neuroblastoma cells (Fig. 1a and Supplementary Fig. 1). We used total internal reflection fluorescence (TIRF) illumination (Fig. 1b) to perform single-particle tracking photoactivated localization microscopy^29^ (sptPALM) at 50 Hz for a duration of 160s. This allowed visualization of the substrate-attached cell surface comprising of the plasma membrane and its subjacent cytoskeleton with ~100 nm axial resolution (Fig. 1c). This imaging paradigm has provided critical insights into the interactions of intracellular signaling molecules, such as Rac1^30^ and KRas^31^, as well as scaffolding proteins, including caveolins^32^, with the plasma membrane. Capitalizing on this approach, we first captured a diffraction-limited TIRF image of Tau^WT^-mEos3.2 before photoconversion in the green (491 nm) emission channel (Fig. 1d). We then recorded a time series of detections of photoconverted Tau^WT^-mEos3.2 molecules near the plasma membrane in the red (561 nm) emission channel, with a localization precision of ~47 ± 6.5 nm (mean ± s.d.) (Fig. 1e,f and Supplementary Fig. 2). This allowed us to construct 5,247 ± 2,373 and 835 ± 495 (mean ± s.d.) Tau^WT^-mEos3.2 trajectories per cell that lasted at least 8 and 20 consecutive image frames, respectively (Fig. 1e and Supplementary Fig. 3). Given that free cytosolic molecules have a large diffusion coefficient of ~10-100 μm^2^/s^33,34^, they would not last ≥8 consecutive frames under TIRF illumination^35^. Thus, the detected trajectories likely represent Tau molecules trapped by plasma membrane components and the associated cytoskeleton. In line with this notion, only a few free cytosolic mEos3.2 trajectories were detected in the TIRF plane with our imaging conditions (Supplementary Fig. 3).

**Figure 1.**
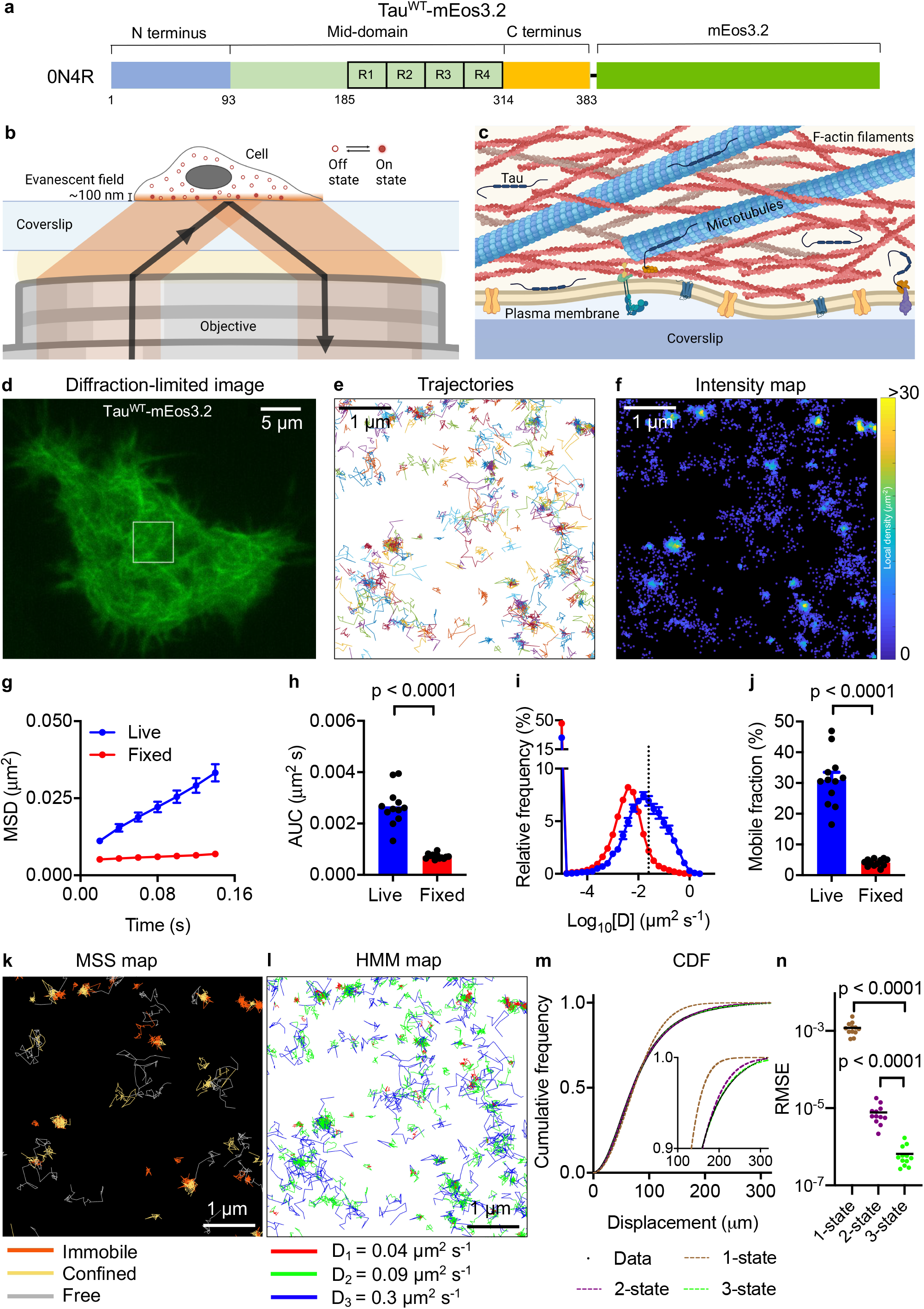
Tau displays heterogenous mobility patterns near the plasma membrane. **a,** Schematic of the most prevalent 0N4R Tau isoform carboxy-terminally tagged with mEos3.2 (Tau^WT^-mEos3.2). **b,** Schematic of detection of Tau^WT^-mEos3.2 molecules in a TIRF microscopy setup. **c,** Schematic of the plasma membrane and subjacent cytoskeleton captured using TIRF imaging as shown in panel **b**. **d,** Representative diffraction-limited TIRF image of an N2a cell expressing Tau^WT^-mEos3.2 acquired in the green emission channel before sptPALM imaging. **e,f,** Maps of trajectories (**e**) and intensities **(f)** of Tau^WT^-mEos3.2 molecules corresponding to the boxed region highlighted in D. The color bar in **f** indicates the local density of each detection computed within a radius of 30 nm. **g-j,** Comparison of Tau^WT^-mEos3.2 mobility parameters in live and fixed cells. **g,** Average MSD as a function of time and **h,** the corresponding area under the curve. **i,** The distribution of the diffusion coefficients in a semi-log plot and **j,** the corresponding mobile fraction. **k,** A map of trajectories annotated as immobile, confined and free state using MSS analysis. **l,** A map of trajectories annotated as distinct diffusive states using the 3-state hidden Markov model. **m, n,** Fits (m) of 1-state (red dashed line), 2-state (magenta dashed line) and 3-state (green dashed line) models to the experimental data (black dotted line) and the root mean square error of all the tested models (n). Error bars, s.e.m. n = 12 live cells and 12 fixed cells. Statistical analysis was performed using the Student’s *t*-test (**h**, **j**) and one-way ANOVA (**n**).

### Tau exhibits heterogeneous mobility pattern near the plasma membrane

We analyzed the mobility pattern of Tau by analyzing its trajectories. Typically, the interaction of a protein with itself or different plasma membrane-associated components manifests as different motion states, ranging from almost stationary to freely diffusive and directed movements^36^. We therefore investigated whether Tau is static or dynamic near the plasma membrane by comparing the mobility of Tau molecules in live and fixed cells. For that, we computed the average mean square displacement (MSD) and the frequency distribution of the instant diffusion coefficient of all trajectories from each analyzed cell (Fig. 1g-j). As expected, the slope of the average MSD curve increased, and the diffusion coefficient distribution was broader and shifted towards higher values in live cells compared to fixed cells, indicating the presence of mobile and immobile pools of Tau molecules near the plasma membrane in live cells (Fig. 1g-j and Supplementary Fig. 4).

We next performed a more detailed quantification of Tau’s complex motion pattern using several complementary approaches (Fig. 1k-n). First, we performed moment scaling spectrum (MSS) analysis, a method widely used to characterize motion states of receptors and transmembrane proteins^37^, and estimated the slope of the MSS of Tau^WT^-mEos3.2 trajectories. We found that near the plasma membrane, Tau existed in three motion states: immobile, confined, and freely diffusive (Fig. 1k). The confinement radii of the immobile and confined states were 66.4 ± 20.2 nm and 88.5 ± 38 nm (mean ± s.d.), respectively (Supplementary Fig. 5a-c). We then analyzed the trajectories using hidden Markov models (HMMs) that assume molecules are switching between discrete diffusive states to estimate the associated model parameters. This analysis revealed that Tau^WT^-mEos3.2 molecules displayed at least three distinct diffusive states, with apparent diffusion coefficients of 0.04 ± 0.01 μm^2^/s, 0.1 ± 0.01 μm^2^/s and 0.29 ± 0.02 μm^2^/s (mean ± s.d.), ranging from immobile to confined to free diffusion (Methods, Fig. 1l, and Supplementary Fig. 5d-f). Finally, we fitted Brownian motion models to the cumulative frequency distribution of the frame-to-frame displacement of Tau^WT^-mEos3.2 molecules and found that the three diffusive state model described the data well (Fig. 1m,n). Together, these results provide evidence that Tau exhibits complex diffusion patterns near the plasma membrane, possibly reflecting Tau’s interactions with each other or its partners in this compartment.

### Tau forms dynamic nanometer-sized hot spots near the plasma membrane

Single-molecule imaging studies have revealed that receptors and signaling molecules form hot spots (a subset of which is known as nanoclusters or nanodomains) at the plasma membrane^27^. We therefore examined whether the observed confined diffusion is due to the similar trapping of Tau in hot spots. Indeed, we found signatures of Tau hot spots with high localization density in the intensity maps that were not detectable in the diffraction-limited TIRF image (Fig. 2a,b). We quantitatively characterized these regions based on both spatial and temporal correlations between individual detections associated with the hot spots as follows. Tessellation-based spatial segmentation algorithms have been widely employed to quantify protein clusters from the sptPALM data^38,39^. Using this approach, we first generated a Voronoï diagram of all localizations, with polygons centered on each single-molecule localization in cells (Fig. 2c). We then identified the cell contour and the potential locations of Tau hot spots based on the normalized localization detection parameters (Methods, Fig. 2d). Next, to characterize their temporal dynamics, we performed time-correlated PALM (tcPALM) analysis, which has previously revealed transient clustering of RNA polymerase^40,41^ and membrane receptors^32^. Here, we computed the time series of molecular detections of individual hot spots and found that the detections were not uniformly distributed but were correlated and clustered in time, indicating the transient nature of these hot spots near the plasma membrane (Fig. 2e). This effect was more apparent in the cumulative detections of molecules, where we observed sudden changes in the slope of detection frequency, marking the duration during which we could detect the hot spots (Fig. 2e). We also found that the trajectories within the hot spots were highly confined within a radius of 71.8 ± 31 nm (mean ± s.d.) (Fig. 2f, Supplementary Fig. 5g). Furthermore, the mobility of Tau^WT^-mEos3.2 molecules in these hot spots was significantly lower than their overall mobility (Fig. 2g-i). To better characterize this confinement, we computed the angles between two successive translocations from three consecutive points and the angular distribution of Tau molecules within the hot spots (Fig. 2j) that together allows for understanding the geometry of the local environment of the molecule^42^. The angular distribution of Tau within the hotspots was shifted towards 180° and significantly different from what is expected from random motion (Fig. 2j) and consistent with motion within a confined space. We next estimated the lifetime of each burst (burst duration) and the number of detections per burst (burst size). The average burst duration and size of the Tau hot spots were 11.9 ± 10.9 s and 136.1 ± 129.2 detection counts (mean ± s.d.), respectively (Fig. 2k,l). Finally, we found that TauWT-mEos3.2 molecules also formed hot spots and displayed heterogeneous mobility patterns in HEK293T cells (Supplementary Fig. 6) similar to those observed in N2a cells, indicating that the dynamic behavior of Tau is conserved across cell types.

**Figure 2.**
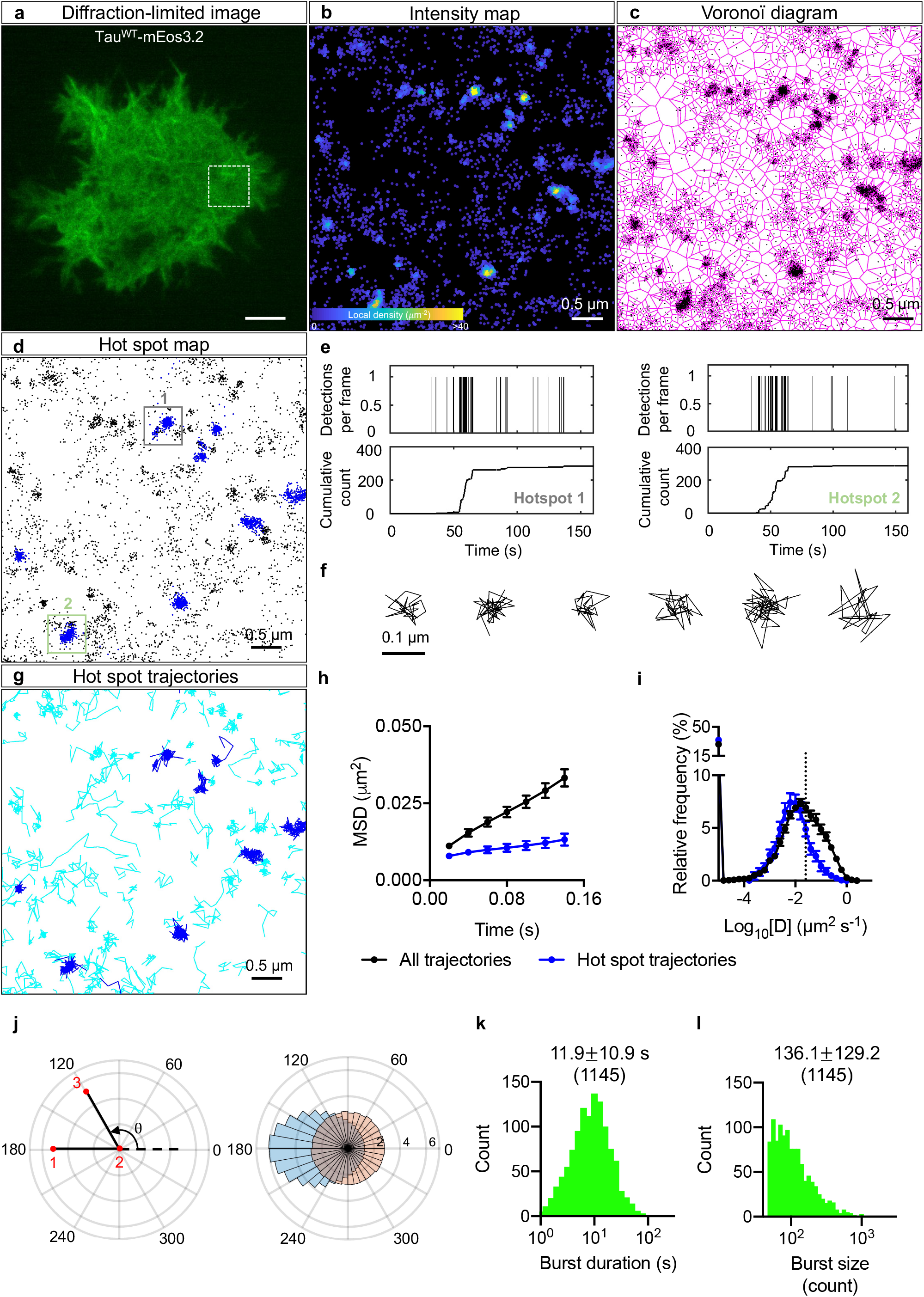
Tau hot spots form transiently near the plasma membrane. **a,** Representative diffraction-limited TIRF image of an N2a cell expressing Tau^WT^-mEos3.2 acquired in the green channel before sptPALM imaging. **b-d,** Intensity map (**b**), Voronoï diagram (**c**), and hot spot map (**d**) of Tau^WT^-mEos3.2 molecules corresponding to the boxed region highlighted in **a**. **e,** Representative time series of detections from two Tau^WT^-mEos3.2 hot spots highlighted in **d**. **f,** Example trajectories detected within hot spots. **g,** Trajectories associated with hot spots are highlighted in blue and the remainder in cyan. **h, i,** Comparison of mobility parameters of TauWT-mEos3.2 molecules associated with hot spots and all the trajectories. Error bars, s.e.m. **j,** Angular distribution of the angle (θ) between the vectors of two successive translocation steps of molecules within hot spots (light blue) compared with that of simulated molecules undergoing Brownian motion (light red). θ is the angle between the vector joining the 1^st^ and 2^nd^ position and the vector connecting the 2^nd^ and 3^rd^ position. **k, l,** The distributions of burst duration (**k**) and size (**l**) of hot spots. The mean ± s.d. are shown together with the number of hot spots analyzed in brackets (n = 12 cells).

### Tau hot spots are not static

To ascertain that the Tau hotspots were not an artifact due to the presence of the mEos3.2 tag, we performed single-molecule imaging of Tau^WT^ fused with the HaloTag in N2a cells incubated with a HaloTag ligand (Janelia Fluor 646). At a 100 pM concentration of the Halo ligand, we only obtained single-molecule signals in N2a cells transfected with Tau^WT^-HaloTag but not in cells transfected with either the free HaloTag or Tau^WT^-FLAG-Tag (Supplementary Fig. 7). By combining Voronoï tessellation-based spatial segmentation and tcPALM analyses, we observed that Tau^WT^-HaloTag molecules also formed hot spots near the plasma membrane (Fig. 3a-e). The average burst duration and size of Tau^WT^-HaloTag hot spots were 10.2 ± 14.2 s and 285.2 ± 519.1 detection counts (mean ± s.d.), respectively (Fig. 3f-h). Membrane protein clusters often exhibit lateral mobility with functional consequences^43^. For example, DC-SIGN and LAT cluster mobility are linked to virus internalization and T cell signaling, respectively^43^. We therefore wondered whether Tau hot spots can also display movements. To test this, we constructed trajectories of Tau^WT^-HaloTag hot spots at 0.2 s intervals over their lifetime (Fig. 3i). We then quantified the MSD curves of hot spot trajectories and found that some hot spots were stationary whereas others were mobile (Fig. 3j). Overall, these results show that Tau forms hot spots that are dynamic at the cell surface.

**Figure 3.**
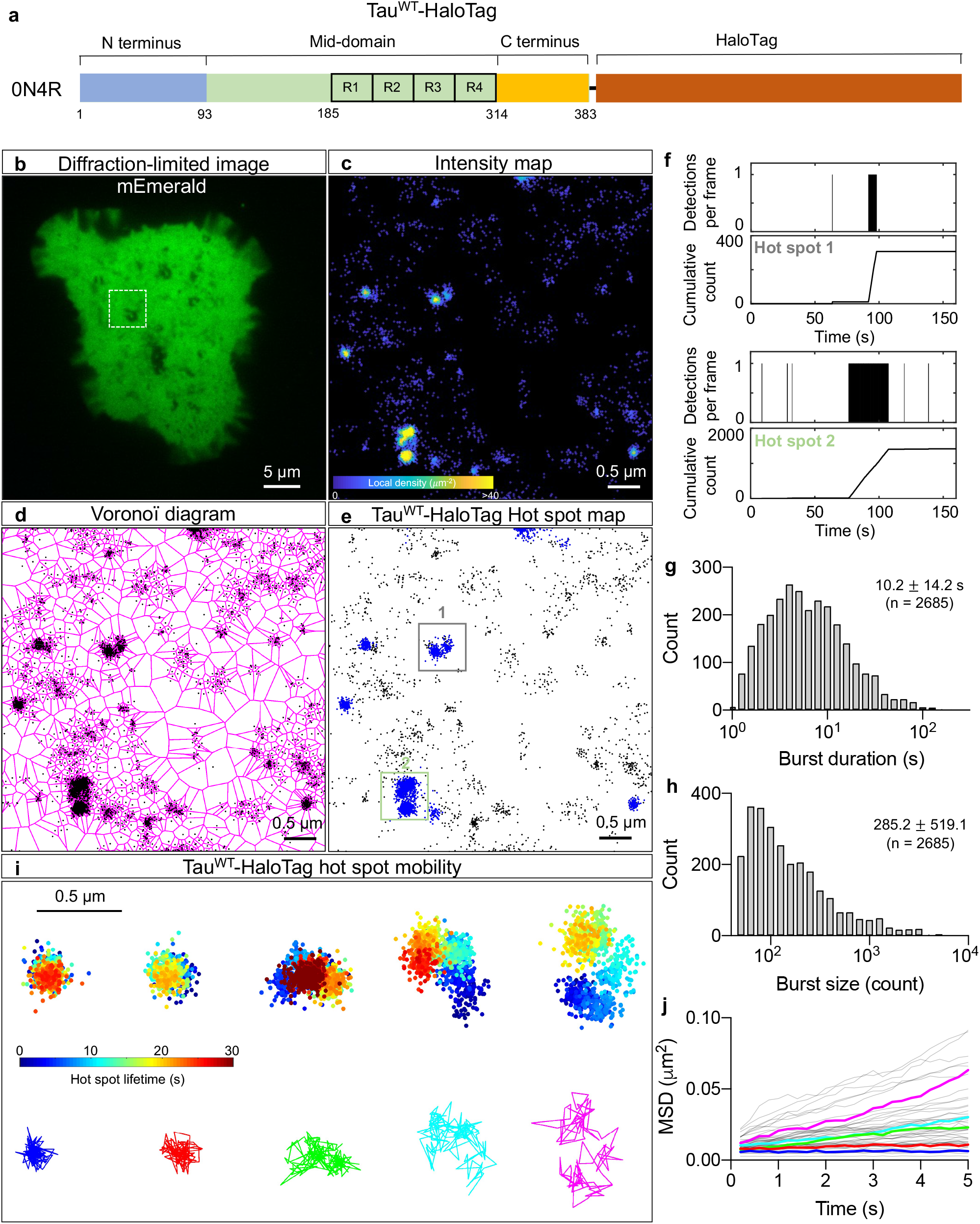
Tau hot spots display distinct motion near the plasma membrane. **a,** Schematic of the 0N4R Tau isoform carboxy-terminally tagged with Halo-Tag (Tau^WT^-HaloTag). **b,** Representative diffraction-limited image of an N2a cell expressing mEmerald acquired in the green emission channel before single-molecule imaging of Tau^WT^-HaloTag in the far-red channel. **c-e,** Intensity map (**c**), Voronoï diagram (**d**), and hot spot map (**e**) of Tau^WT^-HaloTag molecules corresponding to the boxed region highlighted in a. **f,** Representative time series of detections from Tau^WT^-HaloTag hot spots highlighted in d. **g, h,** Distributions of burst duration and size. The mean ± s.d. are shown together with the number of hot spots analyzed in brackets (n = 11 cells). **i,** Examples of hot spots that are immobile (left) and mobile (right). Top, all the localization of hot spots color-coded based on time since hot spot detection. Bottom, trajectories of hot spots constructed by sampling localization belonging to hot spots at 200 ms interval. **j,** MSDs of Tau^WT^-HaloTag hot spots with colored curves representing hot spots highlighted in **i**.

### Nocodazole-mediated microtubule disruption increases the mobility of Tau molecules near the plasma membrane

Given that microtubules are a major binding partner of Tau and that microtubules localize to and interact with the plasma membrane, we next asked whether the mobility pattern of Tau is affected by the availability of microtubules. Indeed, a diffraction-limited TIRF image of the Tau^WT^-mEos3.2 expressing cells readily revealed microtubule-like structures indicative of the association of Tau with the microtubules (Fig. 4a). To assess the effect that microtubules have on Tau’s mobility, we treated the N2a cells with either 5 μM of the microtubule-destabilizing agent nocodazole or DMSO (control) for up to 4 hours. As expected, in the diffraction-limited TIRF images, the density of microtubule filament-like structures visualized by Tau^WT^-mEos3.2 molecules was markedly reduced in nocodazole-treated compared to DMSO-treated live cells (Fig. 4a,b and Supplementary Fig. 8a). A similar effect was observed when imaging the same cell before and after nocodazole treatment (Supplementary Fig. 8b). We also saw a similar decrease in microtubule filaments in fixed, nocodazole-treated N2a cells stained with an anti-tubulin antibody (Supplementary Fig. 9). Consistent with what has been previously reported^44^, we did observe a few nocodazole-resistant microtubules. Remarkably, the trajectory maps showed that Tau molecules could be detected in both DMSO-and nocodazole-treated cells (Fig. 4c,d). We further found that the mobility of Tau was higher in the nocodazole-treated compared to control cells, as assessed by changes in the average MSD (Fig. 4e), the area under the curve (Fig. 4f), the frequency distribution of diffusion coefficients (Fig. 4g), and the percentage of the mobile fraction (Fig. 4h).

**Figure 4.**
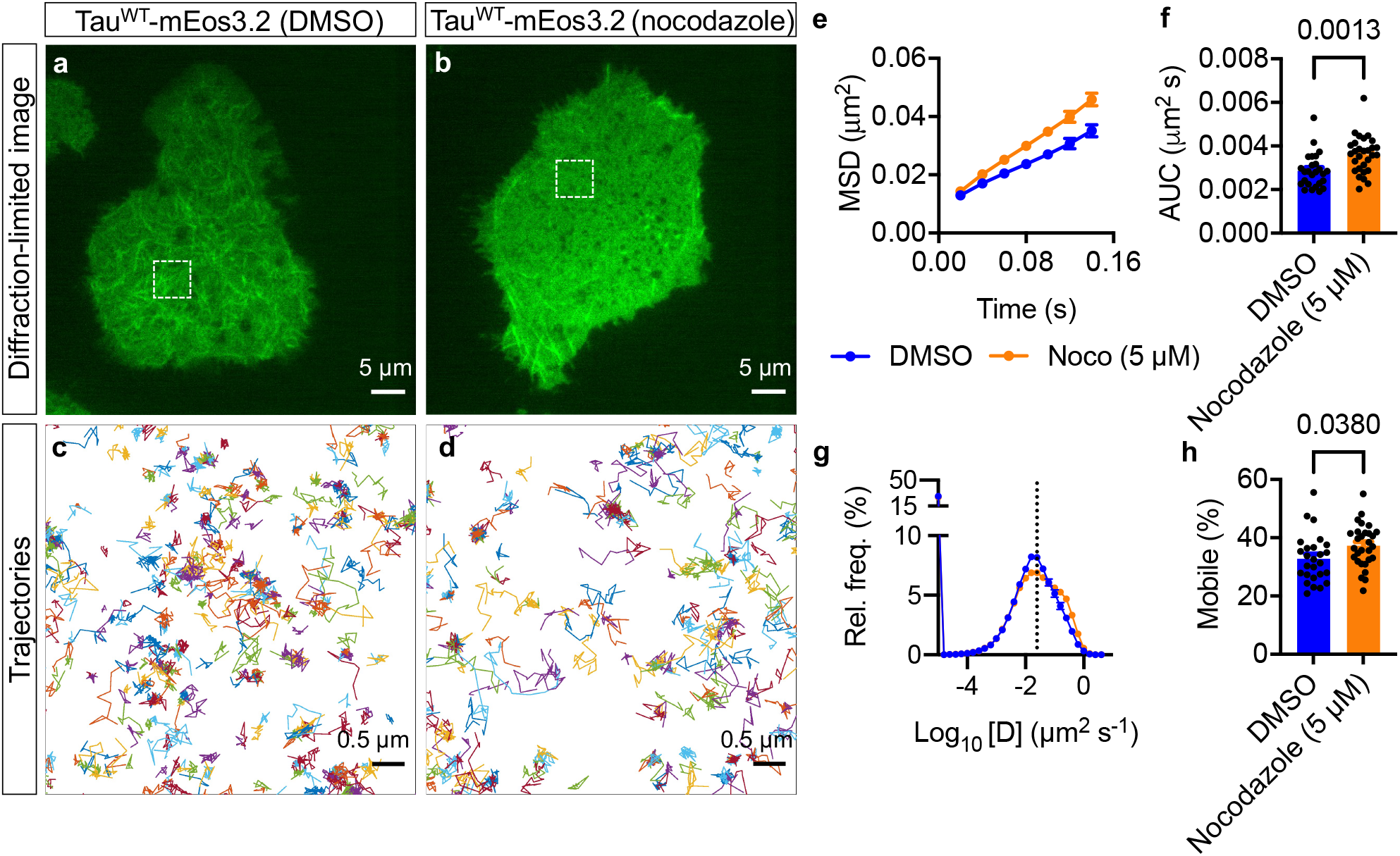
Preventing Tau-microtubule interactions pharmacologically increases Tau’s mobility near the plasma membrane. **a, b,** Representative diffraction-limited TIRF image of N2a cells expressing Tau^WT^-mEos3.2 treated with either DMSO (**a**) or 5 µM nocodazole (**b**) acquired before sptPALM imaging. Scale bar, 5 μm. **c, d,** Maps of trajectories of Tau^WT^-mEos3.2 molecules corresponding to the boxed region highlighted in **a** and **b**. **e-h,** Comparison of Tau^WT^-mEos3.2 mobility parameters in DMSO- and nocodazole-treated live cells. **e,f,** Average MSD as a function of time (**e**) and the corresponding area under the curve (**f**). **g, h,** The distribution of diffusion coefficient shown in semi-log plot and the corresponding mobile fractions (**h**). n = 26 DMSO-treated and 29 nocodazole-treated cells from 3 independent experiments. n = 32 cells expressing Tau^WT^-mEos3.2 and 39 cells expressing Tau^S262E/S356E^-mEos3.2 from 4 independent experiments. Statistical analysis was performed using the Student’s *t*-test.

### Microtubule-binding deficient mutant Tau has increased mobility near the plasma membrane

To further validate this effect, we generated a pseudophosphorylated mutant Tau tagged with mEos3.2 by introducing two serine to glutamate mutations at positions 262 and 356 in the microtubule-binding domain (Tau^S262E/S356E^-mEos3.2) that reduce Tau’s binding affinity to microtubules^45^ (Fig. 5a). As expected, the microtubule filament-like structures were barely visible in Tau^S262E/S356E^-mEos3.2-expressing cells compared to those expressing Tau^WT^-mEos3.2 (Fig. 5b,c and Supplementary Fig. 10), indicating a reduced association of the mutant Tau with the microtubules. Yet, we could readily detect and track a similar number of molecules in both Tau^WT^-mEos3.2-and Tau^S262E/S356E^-mEos3.2-expressing cells (Fig. 5d-e and Supplementary Fig. 11a). Consistent with our nocodazole experiments, we found that Tau^S262E/S356E^-mEos3.2 had higher mobility than Tau^WT^-mEos3.2 (Fig. 5f-i). For both conditions, HMM analysis revealed that Tau molecules displayed at least three distinct diffusive states (Supplementary Fig. 11). Together, these results indicate that the mobility of Tau increases when it is released from microtubules and that the protein exhibits heterogeneous mobility patterns even when the Tau/microtubule interactions are prevented.

**Figure 5.**
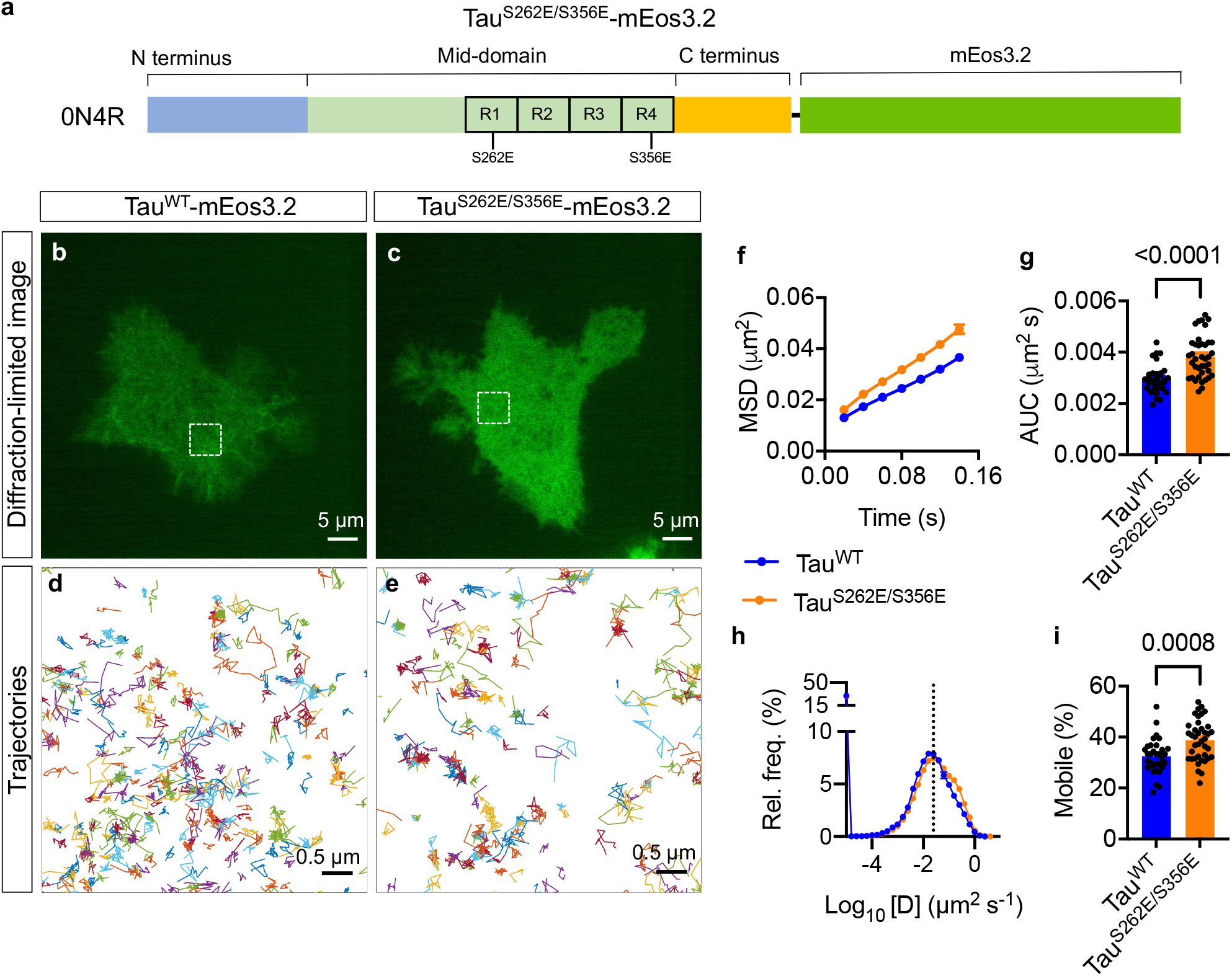
Preventing Tau-microtubule interactions biochemically increases Tau’s mobility near the plasma membrane. **a,** Schematic of 0N4R Tau tagged with mEos3.2 at the C-terminus with S262E/S356E mutations to disrupt microtubule binding. **b, c,** Representative diffraction-limited TIRF image of N2a cell expressing Tau^WT^-mEos3.2 (**b**) and Tau^S262E/S356E^-mEos3.2 (**c**). **d, e,** Maps of trajectories of Tau^WT^-mEos3.2 and Tau^S262E/S356E^-mEos3.2 molecules corresponding to the box region highlighted in **l** and **m**, respectively. **f-i,** Comparison of Tau^WT^-mEos3.2 and Tau^S262E/S356E^-mEos3.2 mobility parameters in live cells. **f, g,** Average MSD as a function of time (**f**) and corresponding AUC (**g**). **h, i,** The distribution of diffusion coefficient (**h**) shown in semi-log plot and the corresponding immobile fraction (**i**). n = 32 cells expressing Tau^WT^-mEos3.2 and 39 cells expressing Tau^S262E/S356E^-mEos3.2 from 4 independent experiments. Statistical analysis was performed using the Student’s *t*-test.

### Tau hot spots are resistant to microtubule perturbations and cholesterol depletion

We next investigated the effect of microtubule perturbations on the dynamics of Tau hot spots. We were able to detect transient hot spots of Tau^WT^-mEos3.2 in nocodazole-treated cells (Fig. 6a-g) and of Tau^S262E/S356E^-mEos3.2 in untreated cells (Fig. 6h-n). In microtubule-perturbed conditions, the burst duration and size of the Tau hot spots were ~13-15 s and ~160-180 detection counts, respectively, and the density of the hot spots was ~0.4-0.5 spots/μm^2^, comparable to the values obtained for Tau^WT^-mEos3.2 in untreated cells. Previous studies have described cholesterol-dependent and −independent mechanisms of protein clustering at the plasma membrane^46^, and cholesterol depletion has been shown to decrease the unconventional release of Tau through the plasma membrane in N2a cells^13^. We therefore examined the effect of cholesterol depletion on the dynamics of Tau hot spots by treating Tau^S262E/S356E^-mEos3.2-expressing N2a cells with 6 mM methyl-β-cyclodextrin (MβCD), a concentration known to strongly deplete cholesterol. Interestingly, there was no apparent effect of cholesterol depletion on the dynamics of Tau hot spots. The burst duration, burst size and density of Tau hot spots were 13.6 ± 14.6 s, 172.9 ± 338.2 detection counts, and 0.46 ± 0.22 spots/μm^2^ (mean ± s.d.), respectively (Fig. 6o-u). Together, these findings that Tau spots are resistant to microtubule perturbations and cholesterol depletion.

**Figure 6.**
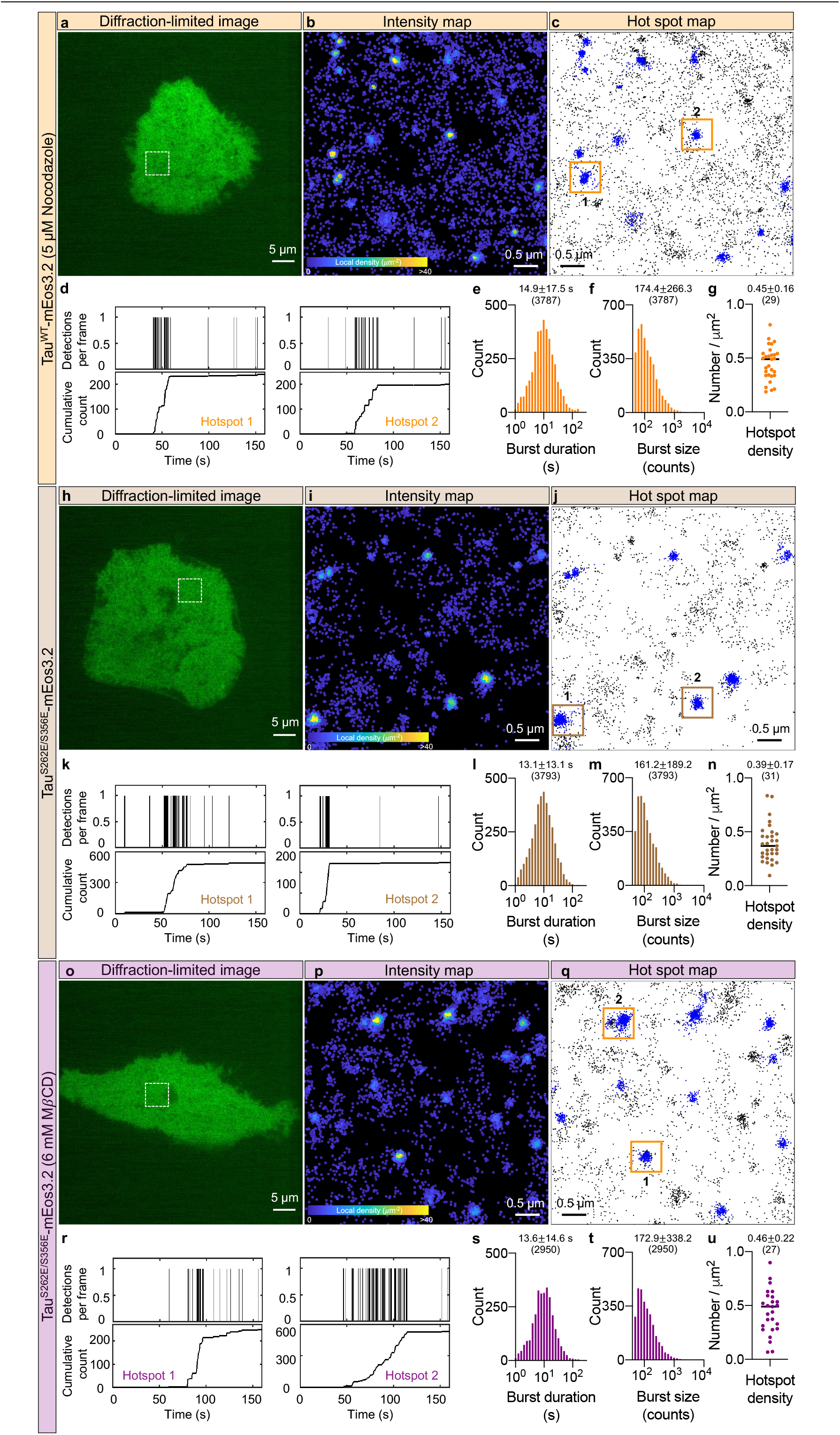
Tau hot spots near the plasma membrane are resistant to both microtubule perturbation and cholesterol depletion. **a,** Representative diffraction-limited TIRF image of an N2a cell expressing Tau^WT^-mEos3.2 in nocodazole-treated live cells. **b, c,** Maps of intensities (**b**) and hot spots (**c**) of Tau^WT^-mEos3.2 corresponding to the boxed region highlighted in **a**. **d,** Representative time series of detections from Tau^WT^-mEos3.2 hot spots highlighted in **d**. **e, f,** Distribution of burst duration and size. **g,** Variations in the density of detected hot spots across cells. n = 29 nocodazole-treated cells from 3 independent experiments. **h,** Representative diffraction-limited TIRF image of an N2a cell expressing Tau^S262E/S356E^-mEos3.2 in live untreated cells. Scale bar, 5 μm. **i, j,** Maps of intensities (**i**) and hot spots (**j**) of Tau^S262E/S356E^-mEos3.2 corresponding to the boxed region highlighted in **h**. **k,** Representative time series of detections from Tau^WT^-mEos3.2 hot spots highlighted in **h**. **l, m,** Distributions of burst duration and burst size. **n,** Variations in the density of detected hot spots across cells. n = 39 cells expressing Tau^S262E/S356E^-mEos3.2 from 4 independent experiments. **o,** Representative diffraction-limited TIRF image of an N2a cell expressing Tau^S262E/S356E^-mEos3.2 in live cholesterol-depleted cells. **p, q,** Maps of intensities (**p**) and hot spots (**q**) of Tau^S262E/S356E^-mEos3.2 corresponding to the boxed region highlighted in **o**. **r,** Representative time series of detections from Tau^WT^-mEos3.2 hot spots highlighted in **q**. **s, t,** The distributions of burst duration and burst size. **u,** Variations in the density of detected hot spots across cells. n = 27 cells expressing Tau^S262E/S356E^-mEos3.2 treated with 6 mM MβCD from 4 independent experiments. In **e, f, l, m, s, t,** The mean ± s.d. are shown together with the number of hot spots analyzed in brackets. In **g, n, u,** The mean ± s.d. are shown together with the number of cells analyzed in brackets.

## DISCUSSION

The cytosolic leaflet of the plasma membrane has long been implicated as a site where Tau executes several of its critical physiological and pathological functions. Yet, due to the limitations of conventional cell biological techniques, our understanding of this protein’s behavior and organization near the plasma membrane of live cells remains scarce. In this study, we describe how Tau diffuses and organizes near the plasma membrane at a single-molecule level resolution. Specifically, we identified distinct kinetic subpopulations of Tau in this compartment. We further found that Tau formed hot spots of diameter ~140 nm, displaying stationary and mobile phases. Moreover, these hot spots lasted tens of seconds and were resistant to microtubule perturbations and cholesterol depletion. Together, our study provides fundamental new insight into the complex spatiotemporal organization of Tau near the plasma membrane.

What are these Tau hot spots? Previous studies have reported liquid-liquid phase separation of Tau, causing the formation of Tau condensates in solutions^47^ and the cytosol of live cells^48^. Moreover, in a cell-free system, microtubules have been shown to catalyze Tau condensation and concentrate Tau molecules in focal regions^49,50^. In these studies, Tau condensates in the cytoplasm and on the microtubules were detectable using conventional diffraction-limited techniques. In contrast, in our cellular system, we did not observe these condensate-like structures near the plasma membrane; our diffraction-limited TIRF images showed either a uniform or microtubule filament-like organization of Tau in this compartment. Importantly, we detected Tau hot spots whose sizes were smaller than the diffraction limit (<250 nm) exclusively in single-molecule localization microscopy images. In addition, inhibiting Tau/microtubule interactions, using either microtubule destabilizing agent or microtubule binding-deficient mutant, did not affect the dynamics of the hot spots. These observations together suggest that these hot spots may differ from the previously reported Tau condensates. Tau phase separation has been suggested as an intermediate step in the Tau aggregation process. It is tempting to speculate that Tau hot spots and condensates are different stages of Tau aggregation pathways. Whether Tau hot spots and condensates interconvert or represent intermediates in divergent Tau aggregation pathways, some leading to fibrils and others not, remains to be investigated.

How do these hot spots form? Protein clustering has emerged as a dominant feature of cellular organization^27,51^. Interestingly, the dynamics of Tau hot spots near the plasma membrane resembled that of membrane receptor clusters^32^ and RNA polymerase II clusters^40,52^. Protein clusters can form either through self-association or interactions with their binding partners. Therefore, one possibility is that the preexisting clusters of Tau’s interaction partner(s) such as Fyn and annexin may serve as sites to capture and trap Tau molecules as they pass by. Alternatively, Tau may itself form assemblies near the plasma membrane, influencing signaling pathways in the local environment. A possible next step forward would be to characterize the local environment (such as lipid composition) that influences or is influenced by Tau hot spots.

Another interesting observation of our study is that the burst duration (apparent lifetime) of hot spots varied by up to two orders of magnitude for all tested conditions, indicating that there is substantial variability in the kinetics of hot spots and also that multiple mechanisms may determine hot spot formation and stability. Remarkably, the lifetime of a hot spot by far exceeds (by two orders of magnitude) the reported ~40 ms dwell time of Tau on microtubules^53^, hinting that the interactions of Tau with different partners lead to vastly different Tau behaviors and potentially different biological outcomes. It is tempting to speculate that it is not only the dynamics of individual hot spots but also the coordination of multiple hot spots at the plasma membrane that determines Tau’s membrane functions. Whether Tau hot spots contribute to membrane insertion and fibrillization in pathological conditions such as AD remains to be determined.

Protein motion types have been previously linked to conformational changes, protein-protein interactions, and post-translation modifications. Here, we show that Tau, a protein undergoing massive post-translation modifications, exhibits distinct motion states given that it undergoes free diffusion along the plasma membrane, it then gets immobilized and trapped inside hot spots, and microtubules could bind and transiently confine it. In our study, the diffusion coefficient of Tau varied over several orders of magnitude suggesting that multiple mechanisms might underlie the heterogenous mobility patterns of Tau near the plasma membrane. Given that a significant fraction of Tau molecules diffuses freely near the plasma membrane, we speculate that free diffusion is a strategy that allows Tau to explore the membrane at high efficiency, and that this pool serves as a reservoir to supply Tau to its multiple interactors at the plasma membrane. Consistent with this notion, blocking Tau’s interactions with its partner microtubules increased the mobile reservoir pool of Tau available for other interactions. This dynamic property of Tau likely allows the protein to transiently interact with multiple partners and thereby execute a rich repertoire of cellular functions in space and time^16,54^.

In summary, our study sheds new light on the spatiotemporal organization of Tau near the plasma membrane. Given Tau’s emerging role as a multifunctional protein in physiology and pathology, we provide a framework to investigate the functions of Tau at the plasma membrane as well as in other cellular compartments.

## ACKNOWLEDGMENTS

We thank Rowan Tweedale for critical reading of the manuscript. We acknowledge Corey Butler and Adel Kechkar for their contributions in the development of PALM-Tracer. The imaging was performed at the Queensland Brain Institute Advanced Microscopy Facility, supported by the Australian Government through the Australian Research Council LIEF grant (LE130100078). We acknowledge support by the Estate of Dr. Clem Jones, the State Government of Queensland (DSITI, Department of Science, Information Technology and Innovation), and the National Health and Medical Research Council of Australia (GNT1176326, GNT1147569, and GA39196) to J.G.

## METHODS

### Cell culture and transfection

N2a mouse neuroblastoma cells were maintained in Dulbecco’s minimum essential medium (ThermoFisher; Catalog No. 11965-092) supplemented with 10% foetal bovine serum (Bovogen; Catalog No. SFBS-F) and 50 U/ml penicillin/streptomycin (Gibco; Catalog No. 15140-122). Cells were grown at 37°C in 5% CO_2_. 250,000 cells per well were plated in 12-well plates for 18 h before transfection. Lipofectamine^TM^ LTX and Plus reagent (Invitrogen; Catalog No. 15338030) were used for cell transfections as per the manufacturer’s instructions. For imaging, the cells were dissociated with Trypsin-EDTA (0.25%; Catalog No. 25200-056), centrifuged, and ~32,000 cells were replated on poly-D-lysine-coated 35 mm glass-bottom culture dishes (Ibidi) for one hour. The media was then full changed, and cells were imaged after 16 to 40 h.

### Plasmids

Mammalian expression plasmids were made using human tau cDNA for isoform 0N4R and the human cytomegalovirus (CMV) promoter. Fusion proteins were created with a C-terminal tag of mEos3.2, HaloTag or FLAG-tag (DYKDDDDK). The plasmid mEos3.2-C1 was a gift from Michael Davidson and Tao Xu (RRID:Addgene_54550)^55^. Pseudo-phosphorylated Tau at Ser262 and Ser356 was made by substitution of serine by glutamic acid. Plasmids for control expression of tagged proteins were created by deletion of the tau encoding sequence.

### Western blot analysis

N2a cells were collected for expression validation from the 12-well plate at 24 h post transfection in RIPA buffer (Cell Signalling Technologies; Catalog No. 9806) in the presence of phosphatase inhibitor (Roche; Catalog No. 04906837001) and protease inhibitor (Roche; Catalog No. 04693159001). Samples were diluted in Laemmli buffer, sonicated and heated to 95°C for 10 minutes, then separated using SDS-PAGE on 4-20% Criterion TGX (BioRad; Catalog Nos. 5671084 and 5671085) gradient gels at 250 V. The samples were transferred to nitrocellulose membranes (Merck; Catalog No. HATF00010) for 50 minutes at 400 mA. The membranes were blocked in TBS containing Odyssey Blocking Buffer (LI-COR; Catalog No.927-50000) and incubated in primary antibodies (Tau5 1:5,000) overnight at 4°C followed by incubation with the secondary antibody for one hour at RT. Membranes were imaged on the LI-COR Odyssey scanner using the Image Studio software (LI-COR; Catalog Nos. 926-32211 and 926-68070).

### TIRF microscopy and single-molecule imaging

For live-cell TIRF microscopy, transfected cells were imaged using an iLas^2^ azimuthal TIRF illumination system (Roper Scientific) mounted on a Nikon Ti-E inverted microscope, with a 100x/1.49 NA oil-immersion TIRF objective (CFI Apochromat, Nikon) and Evolve 512 Delta EMCCD cameras (Teledyne Photometrics). Image acquisition was performed using MetaMorph (version 7.10.1.161, Molecular Devices). Single-molecule imaging of Tau tagged with mEos3.2 was performed at 50 Hz to record 8,000 frames per cell at 37 °C. Cells were washed and incubated with buffer A (145 mM NaCl, 5 mM KCl, 1.2 mM Na2HPO4, 10 mM D-glucose, 20 mM Hepes, pH 7.4) during imaging. To perform sptPALM, we used a 405 nm laser (Stradus 405, Vortran Laser Technology) to photoconvert the mEos3.2-tagged molecules and a 561 nm laser (Cobolt Jive, Cobolt Lasers) for excitation of the photoconverted molecules. To activate and detect the mEos3.2 signal from the background signals, we used a TIRFM GFP/RFP filter cube (Nikon corporation) in the microscope body, and a T565lpxr long-pass dichroic beam splitter, and an ET600/50m emission filter (Chroma Technology) in the TwinCam (Cairn Research) dual emission image splitter transmission arm. The 405 nm laser power density was between 3.4 x 10^-6^ and 9.5 x 10^-5^ kW/cm^2^, and the 561 nm laser power density was set to ~0.14 kW/cm^2^. To track Tau^WT^-HaloTag, we first incubated cells co-expressing Tau^WT^-HaloTag and the cell fill mEmerald with the Halo ligand (2 pM) for 15 min at 37°C. Then, cells were washed and Tau^WT^-HaloTag molecules were tracked using a 642 nm laser at a power density of 0.12 kW/cm^2^, a ZT405/488/561/647rpc quad-band dichroic beam splitter and ZET405/488/561/640m emission filter (Chroma Technology) in the microscope body, and a ZT647rdc dichroic beam splitter and an ET690/50m emission filter (Chroma Technology) in the TwinCam transmission arm.

### Nocodazole and MβCD treatments

For nocodazole treatment experiments, cells were incubated with either nocodazole (5 μM) or DMSO mixed with culture medium. Cells were then washed and incubated with buffer A containing either nocodazole (5 μM) or DMSO during imaging. For MβCD experiments, cells were washed and incubated with buffer A containing 6 mM MβCD. Imaging was performed between 5 and 40 minutes post MβCD treatment.

### SptPALM analysis

#### Tracking

We localized and tracked individual Tau molecules tagged with mEos3.2 as previously described^38,56–58^. Briefly, individual molecules were localized using a wavelet-based segmentation algorithm^59^ and trajectories were computed using a simulated annealing-based tracking algorithm^60^, with the PALM-Tracer tool that operates as a plugin of Metamorph software (Molecular Devices). We used a frame-to-frame particle-linking distance threshold of 318 nm (3 pixels). We visually inspected for misconnections using the InferenceMAP tool in both fixed and live cells^61^. Of note, we observed a small proportion of mobile molecules in fixed samples, potentially due to incomplete molecular immobilization after fixation using paraformaldehyde^62^.

#### Intensity and trajectory maps

Intensity and trajectory maps were constructed using detections lasting at least four and eight consecutive frames, respectively. For intensity maps, the local density of each detection was determined by computing the number of detections with a circle of 30 nm radius using a custom written code.

#### Mean square displacement and diffusion coefficients

We constructed trajectories of detections that lasted at least eight consecutive frames and computed the MSD of each trajectory. The MSD was fitted by the equation *MSD*(τ) = *a* + 4*Dτ*, where *D* is the diffusion coefficient, *a* is the y-intercept and τis the time shift. Cells with at least 1,000 trajectories lasting at least 8 frames were considered for analysis with MSD and diffusion coefficients. The average MSD of all trajectories from each analyzed cell was fitted by the equation *MSD*(τ) = *a* + 4*D_avg_τ* to estimate the average diffusion coefficient *D_avg_* (Supplementary Fig. 12).

#### Moment scaling spectrum

Using divide-and-conquer moment scaling spectrum (DC-MSS) tool^63^, we performed the moment scaling spectrum (MSS) analysis to Tau^WT^-mEos3.2 trajectories that lasted at least 20 frames, estimated the slope of MSS, and categorized the trajectories into different motion states, as described previously^37,63^. Briefly, for every time shift τ, the moments of displacement, μm, were computed for *m*= 1,…, 6. Next, using the relationship μ_m_(τ) = 4*D_m_τ^αm^*, the generalized diffusion coefficient *D_m_* and the exponent *α_m_* are estimated for each *m*. The plot *αm* versus *m* yields the MSS and the slope of the MSS allows categorizing trajectories into different motion types.

#### Hidden Markov modelling

The vibrational Bayes SPT (vbSPT) tool^64^ was used to analyze Tau^WT^-mEos3.2 and Tau^S262E/S356E^-mEos3.2 trajectories. Cells with ≥1000 trajectories were used for this analysis. HMM approach models the trajectories as random transitions between a set of hidden states with different diffusion coefficients. By applying Bayesian model selection to hidden Markov models, the vbSPT analysis infers the number of hidden diffusive states and the associated parameters from the experimental data. Initially, when we allowed a maximum of 10 hidden states, models with 3 or more states provided the best fit (Supplementary Fig. 5). We have therefore presented the parameters estimated by fitting a three-state model to the experimental data (Fig. 1l and Supplementary Figs. 5f, 6j and 11b).

#### Frame-to-frame displacement analysis

We computed the empirical cumulative frequency distribution of the displacements of Tau molecules tagged with mEos3.2 molecules at 20 ms intervals using the tool ECDF in MATLAB. The one-state model is described by 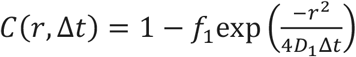, the two-state model by 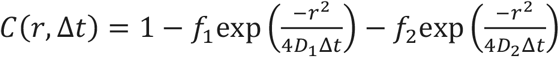, and the three-state model by 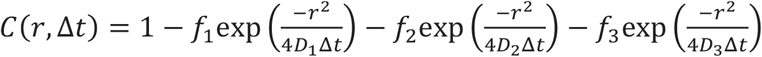. Here, *r* is the displacement, Δ*t* is the time interval (20 ms), *D*_1_, *D*_2_ and *D*_3_ are the diffusion coefficients of the three states, and *f*_1_, *f*_2_ and *f*_3_ are the state occupancies. We fit the predictions of different models to the data using the non-linear regression tool NLINFIT in MATLAB to estimate the model parameters.

#### Voronoï tessellation and time-correlated PALM (tcPALM)

We used a cross-correlation-based method to correct for any motion drift during imaging acquisition^65^. We then used the SR-Tesseler tool^39^ to identify the outline of the cell (object). Tau hot spots were identified as regions with local density at least five-fold greater than the average density of the object. Next, we performed the tcPALM analysis^40,41^ by drawing a square region of interest around each spot and computing the number of detections within the region as a function of time. The start and end points of each burst were identified from the cumulative detections using dark time tolerance of 200 frames. Bursts with at least 50 detections were considered for further analysis. To avoid cell edge confounds due to the folding of membranes at the edges, we analyzed cell areas of ~100-400 μm^2^, excluding cell edges.

#### Statistical analysis

The D’Agostino and Pearson test was used to test for normality, and the Student’s *t*-test was used for statistical comparison when the data were normally distributed. When data sets had more than two groups, an ANOVA was used with appropriate corrections for multiple comparisons. Values are represented as the mean ± s.e.m or mean ± s.d., as indicated in the figure legends. Data were considered significant at p<0.05. GraphPad Prism 9 was used to perform statistical tests and making figures.

