## Supplementary Information for "Tau forms dynamic hot spots that are resistant to microtubule perturbations and cholesterol depletion"

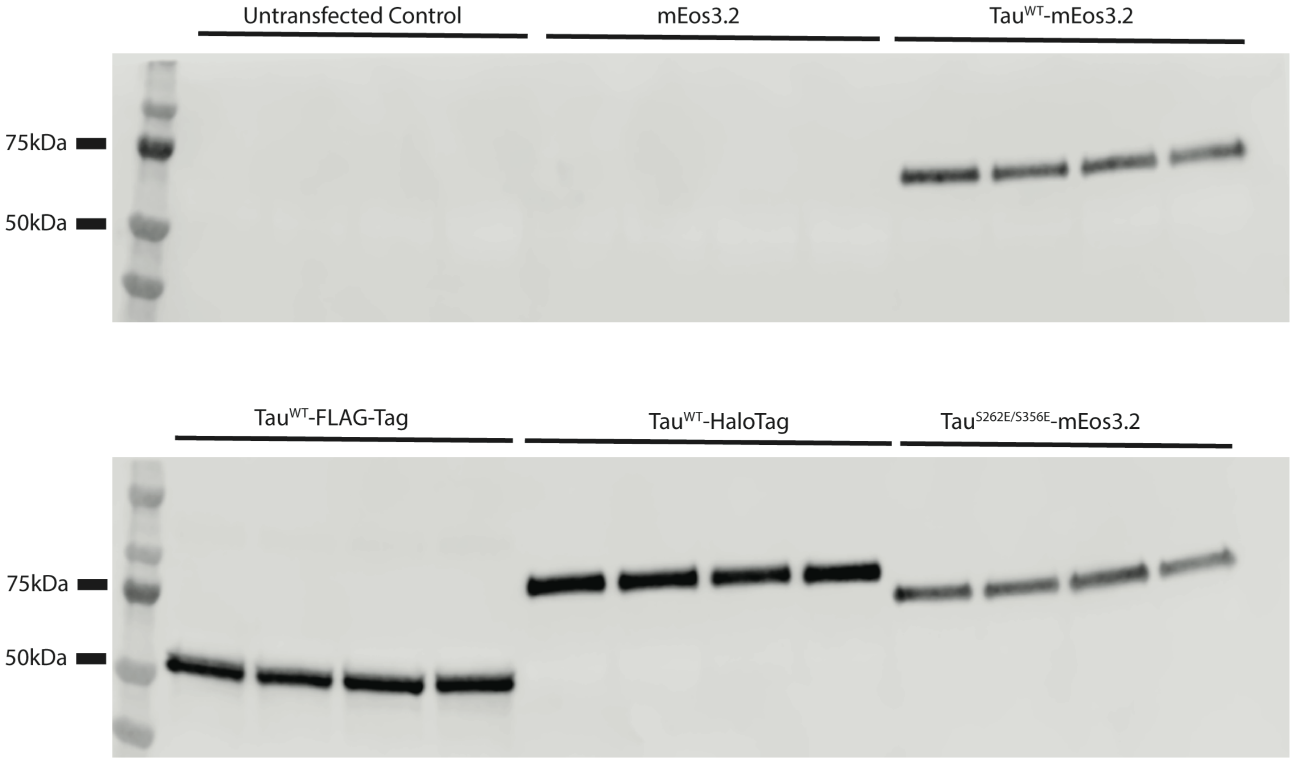


**Supplementary Figure 1. Western blot analysis of expression of different Tau constructs in N2a cells.** Cells transfected with different Tau constructs were analyzed by TAU-5 antibody.

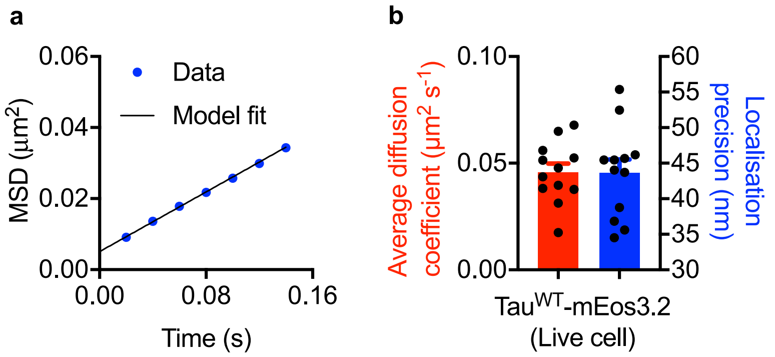


**Supplementary Figure 2. Estimates of the average diffusion coefficients and the localization precision. a,** The average MSD of trajectories from a representative cell was fitted by the equation $MSD(\tau) = 4\sigma^2 + 4D_{avg}\tau$, where $D_{avg}$ is the average diffusion coefficient, $\tau$ is the time lag and $\sigma$ is the localization precision. **b,** Estimates of $D_{avg}$ and $\sigma$ from different cells. $R^2$ was greater than 0.99 for all the fits, and the first four points of the MSD curve were used for fitting.

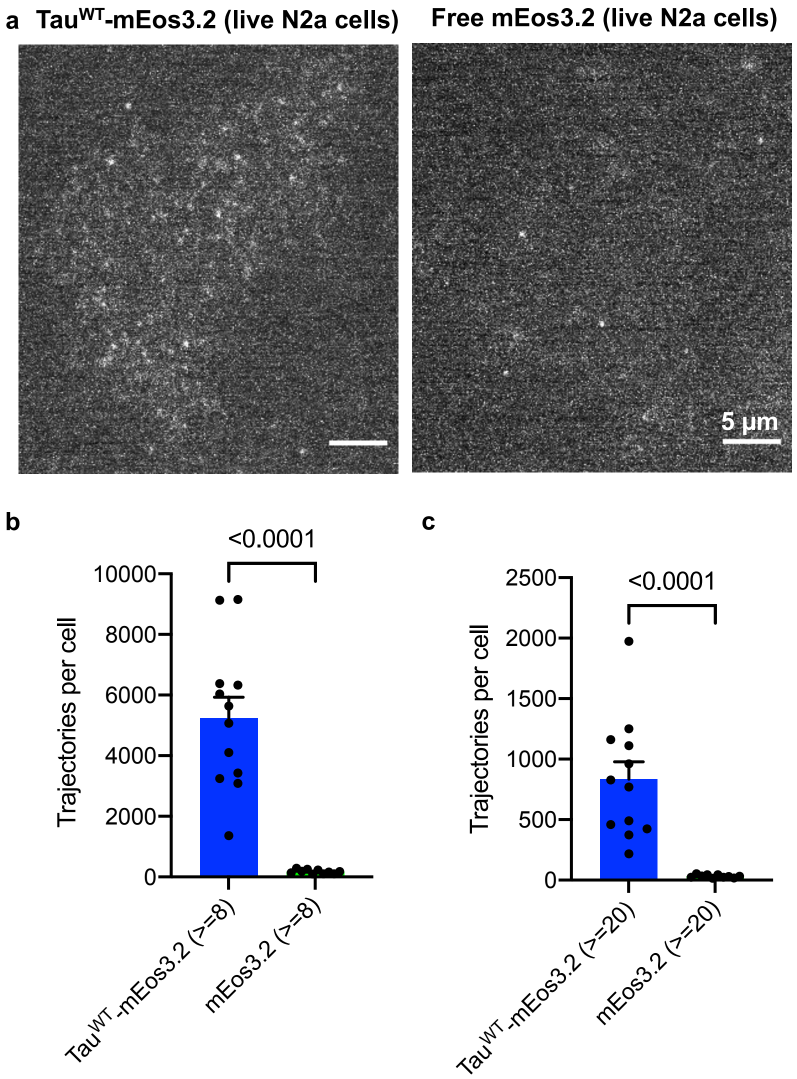


**Supplementary Figure 3. Statistics of trajectories detected in N2a cells expressing Tau$^{WT}$-mEos3.2 or free mEos3.2. a,** Representative images of live N2a cells expressing either Tau$^{WT}$-mEos3.2 (left) or free mEos3.2 (right) showing single molecules detected in one frame. **b,c,** Comparison of detected trajectories lasting at least 8 (**b**) and 20 (**c**) frames in live N2a cells expressing either Tau$^{WT}$-mEos3.2 or free mEos3.2.

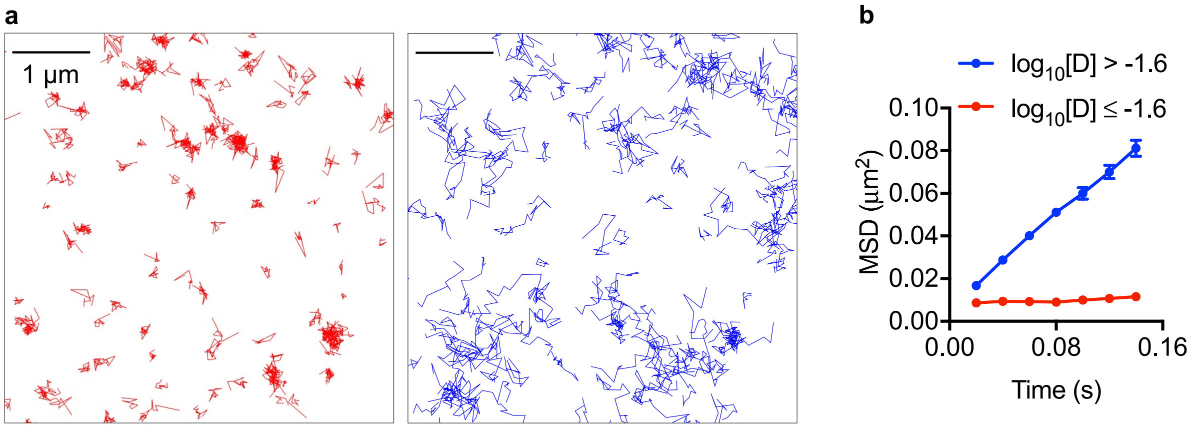


**Supplementary Figure 4. Tau$^{WT}$-mEos3.2 trajectories with varying diffusion coefficients in live N2a cells. a,** Representative region of a N2a cell with Tau$^{WT}$-mEos3.2 having $\log_{10}$ [D] ≤ -1.6 (color-coded in red) and $\log_{10}$ [D] > -1.6 (color-coded in blue). **b,** Average mean square displacement as a function of time for trajectories with $\log_{10}$ [D] ≤ -1.6 and $\log_{10}$ [D] > -1.6 (n = 12 cells).

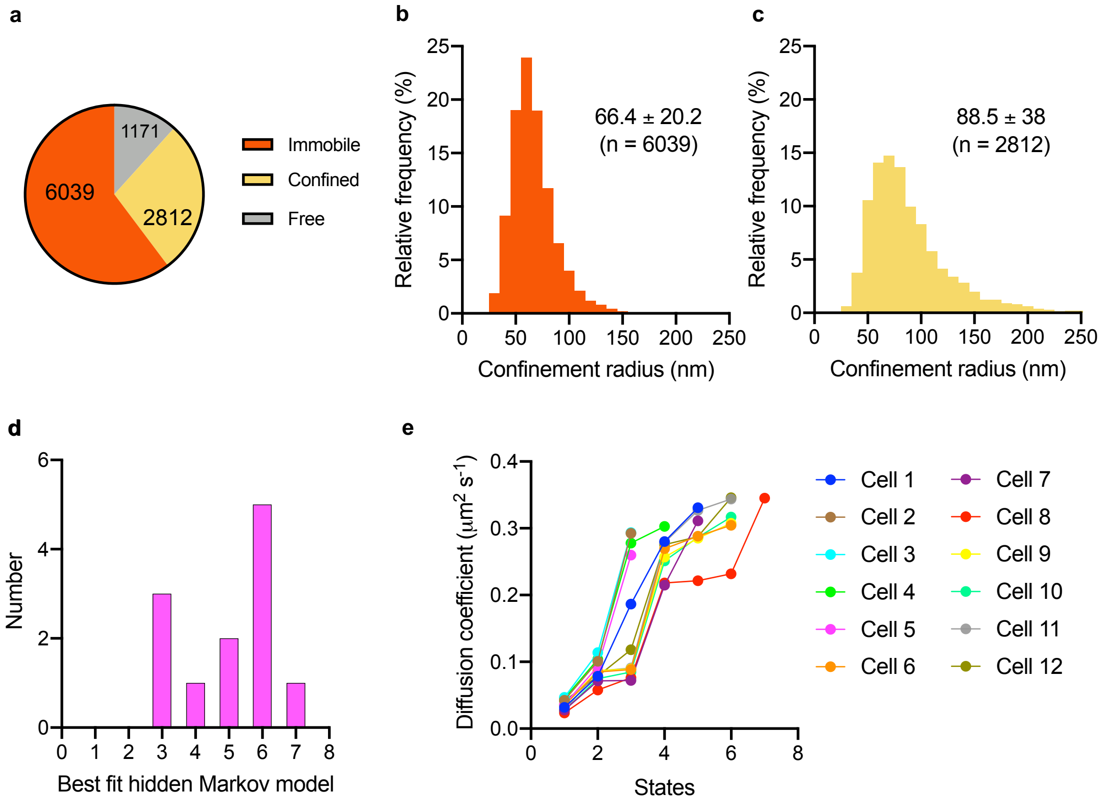


**f**

| | Apparent diffusion coefficient ($\mu m^2\ s^{-1}$) | | State occupancy (%) | |
|---|---|---|---|---|
| | vbSPT | CDF | vbSPT | CDF |
| State 1 | 0.037±0.008 | 0.028±0.003 | 29.8±9.8 | 16.1±9.3 |
| State 2 | 0.095±0.011 | 0.078±0.006 | 45.0±3.2 | 56.2±6.5 |
| State 3 | 0.29±0.02 | 0.029±0.03 | 25.2±10.4 | 27.7±10.8 |

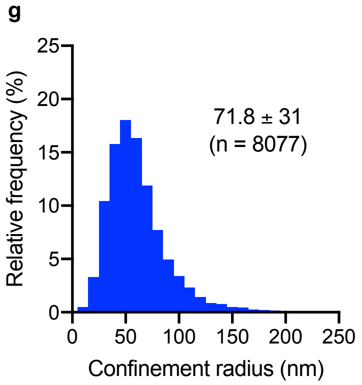


**Supplementary Figure 5. Detailed analysis of Tau$^{WT}$-mEos3.2 trajectories. a,** The relative distribution of trajectories lasting at least 20 frames annotated as immobile, confined, and diffusive fractions using MSS analysis. **b,c,** The confinement radius of immobile (**b**) and confined (**c**) trajectories identified using MSS analysis. **d,** The distribution of the best fit model by allowing a maximum of 10 hidden states during model selection. **e,** The apparent diffusion coefficients of hidden states of the best fit models. **f,** A summary of diffusion coefficients and state occupancy of a three-state model inferred using hidden Markov modeling (vbSPT tool) and CDF fitting. The mean±s.d. are shown. **g,** The confinement radius of trajectories belonging to hot spots. In **b,c,g,** the confinement radius was computed by eigenvalue decomposition of the variance-covariance matrix of particle coordinates. n = 12 cells expressing Tau$^{WT}$-mEos3.2.

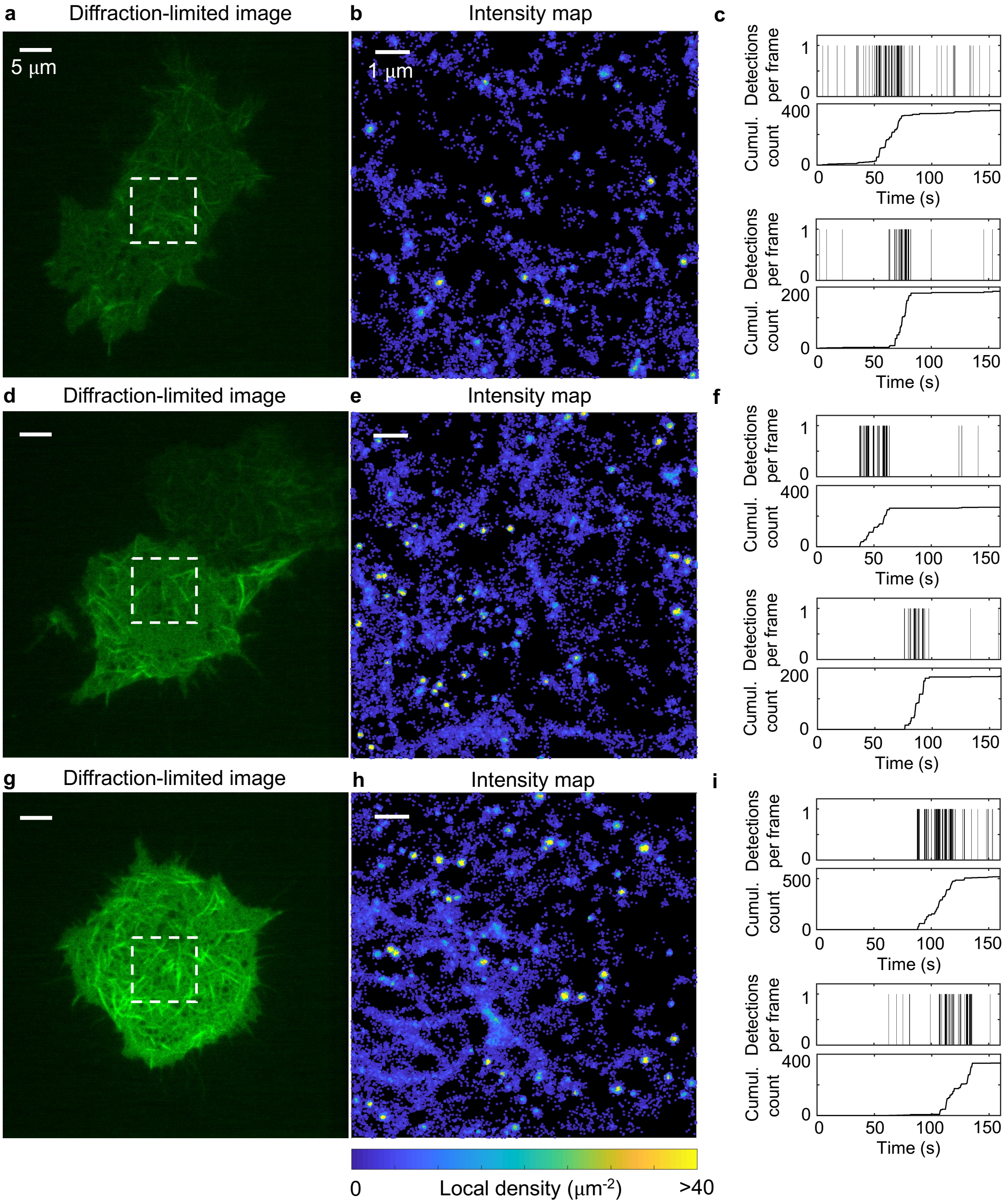


j

| Three-state hidden Markov model (vbSPT tool) | | |
|---|---|---|
| | Apparent diffusion coefficient (µm² s⁻¹) | State occupancy (%) |
| State 1 | 0.037±0.009 | 28.7±10 |
| State 2 | 0.096±0.017 | 46.4±7.5 |
| State 3 | 0.27±0.024 | 24.9±7.3 |

**Supplementary Figure 6. Tau forms hot spots and displays heterogeneous mobility patterns in HEK293T cells. a-i,** Representative low-resolution TIRF images (**a,d,g**) and intensity maps (**b,e,h**) of HEK293T cells expressing $Tau^{WT}$-mEos3.2. Examples of time series of detections from $Tau^{WT}$-mEos3.2 hot spots detected in these cells (**c,f,i**). **j,** A three-state model was the best fit for all cells (n = 9 cells) when a maximum of three states was allowed during model selection. A summary of diffusion coefficients and state occupancy of a three-state model inferred using hidden Markov modeling (vbSPT tool). The mean±s.e.m. are shown.

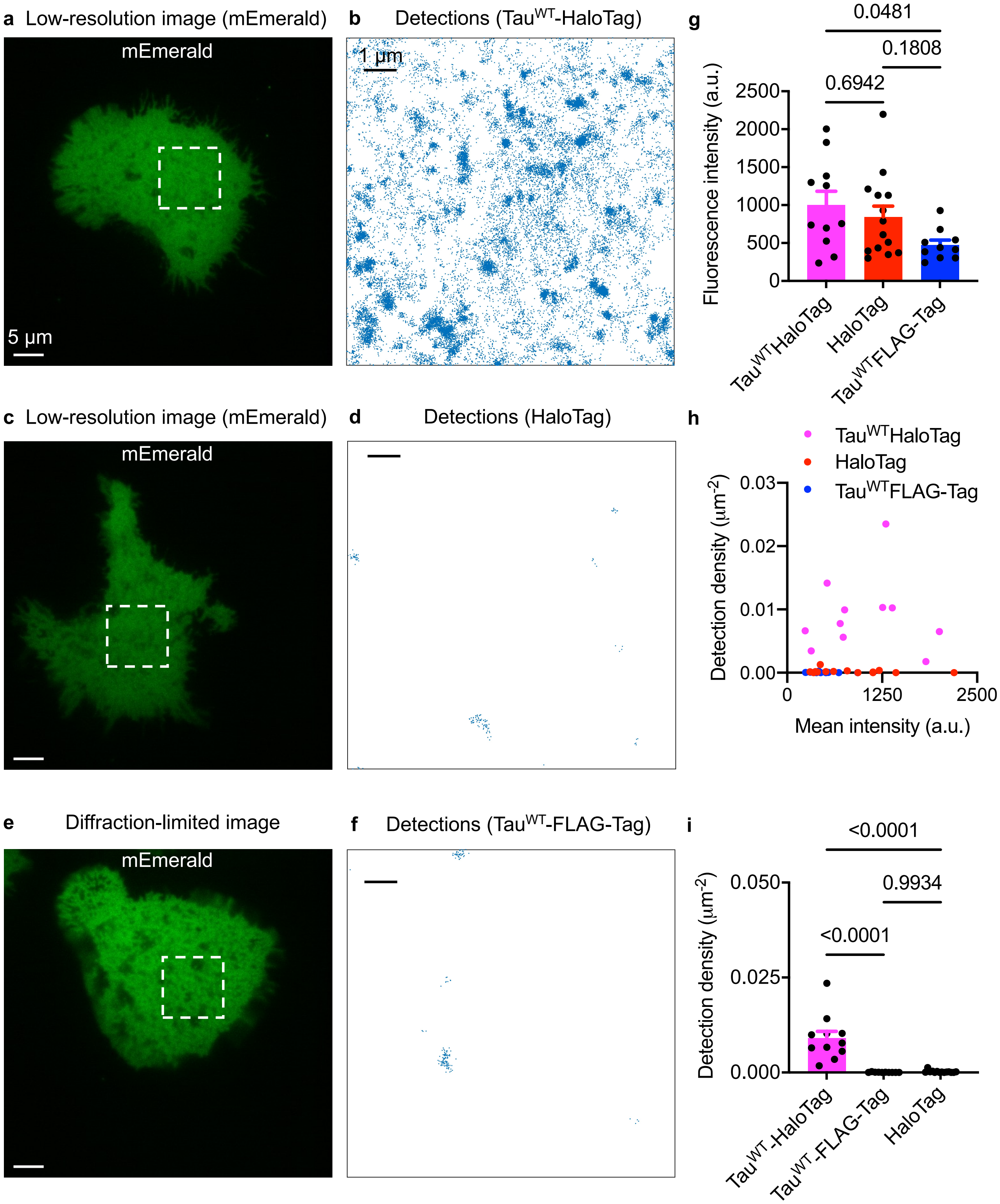


**Supplementary Figure 7. Statistics of detections in N2a cells co-expressing mEmerald and Tau$^{WT}$-HaloTag or Tau$^{WT}$-FLAG-Tag or free HaloTag. a-f,** Representative low-resolution TIRF images of mEmerald (**a,c,e**) and maps of detections lasting at least 4 frames (**b,d,f**) of N2a cells co-expressing mEmerald and Tau$^{WT}$-HaloTag (**a,b**) or free HaloTag (**c,d**) or Tau$^{WT}$-FLAG-Tag (**e,f**). **g,** Comparison of the intensity of mEmerald across conditions. **h,** Correlation between the detection density and the intensity of mEmerald across conditions. **i,** Comparison of the intensity of mEmerald across conditions. The mean±s.e.m. are shown. Statistical analysis was performed using one-way ANOVA.

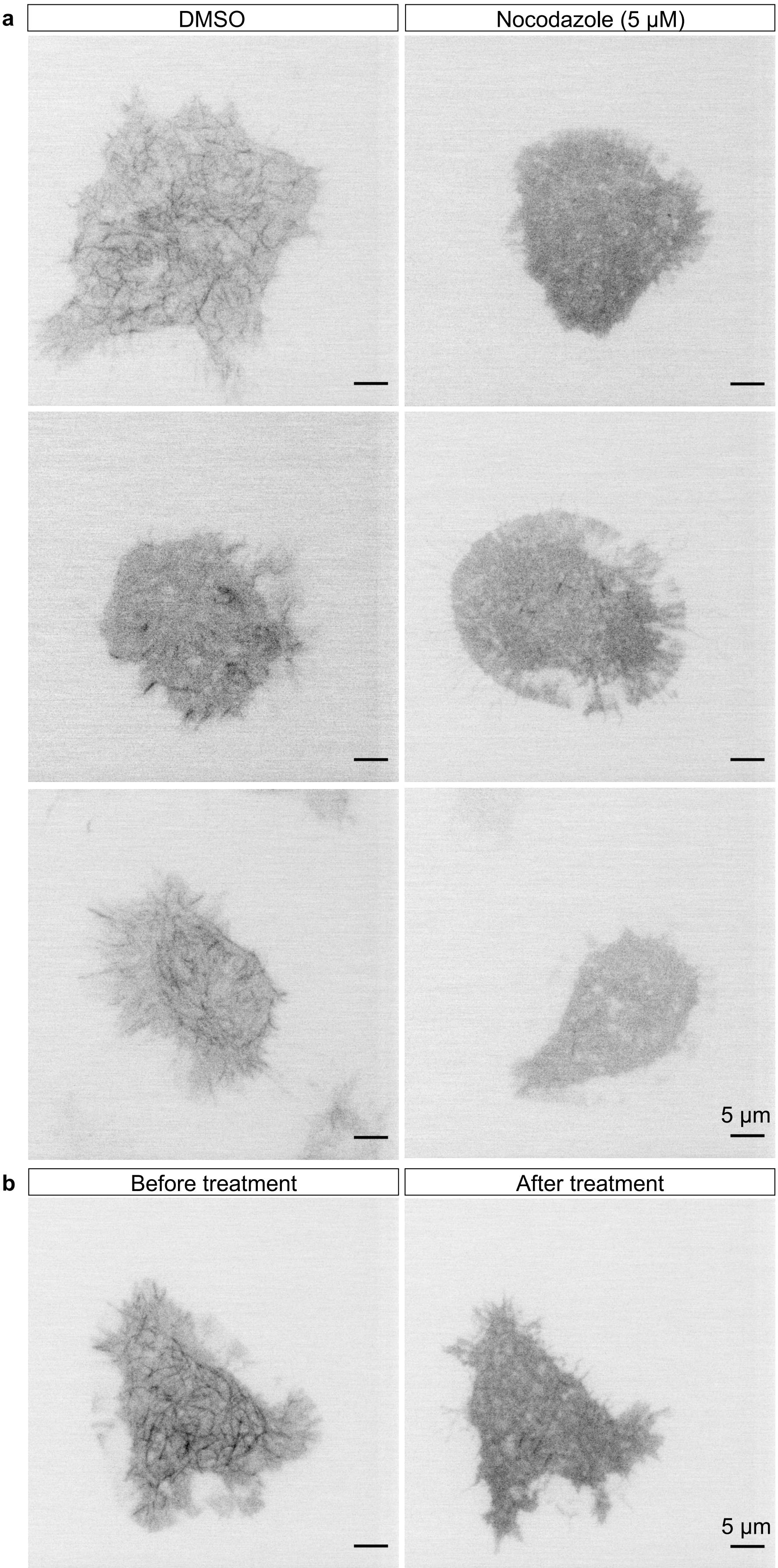


**Supplementary Figure 8. Nocodazole-treated cells have less filament-like structures. a,** Low-resolution TIRF images of $Tau^{WT}$-mEos3.2 in nocodazole-treated (left) and DMSO-treated (right) live cells acquired in the green channel. **b,** Low-resolution TIRF images of $Tau^{WT}$-mEos3.2 in a live cell before and after nocodazole (5 μm) treatment.

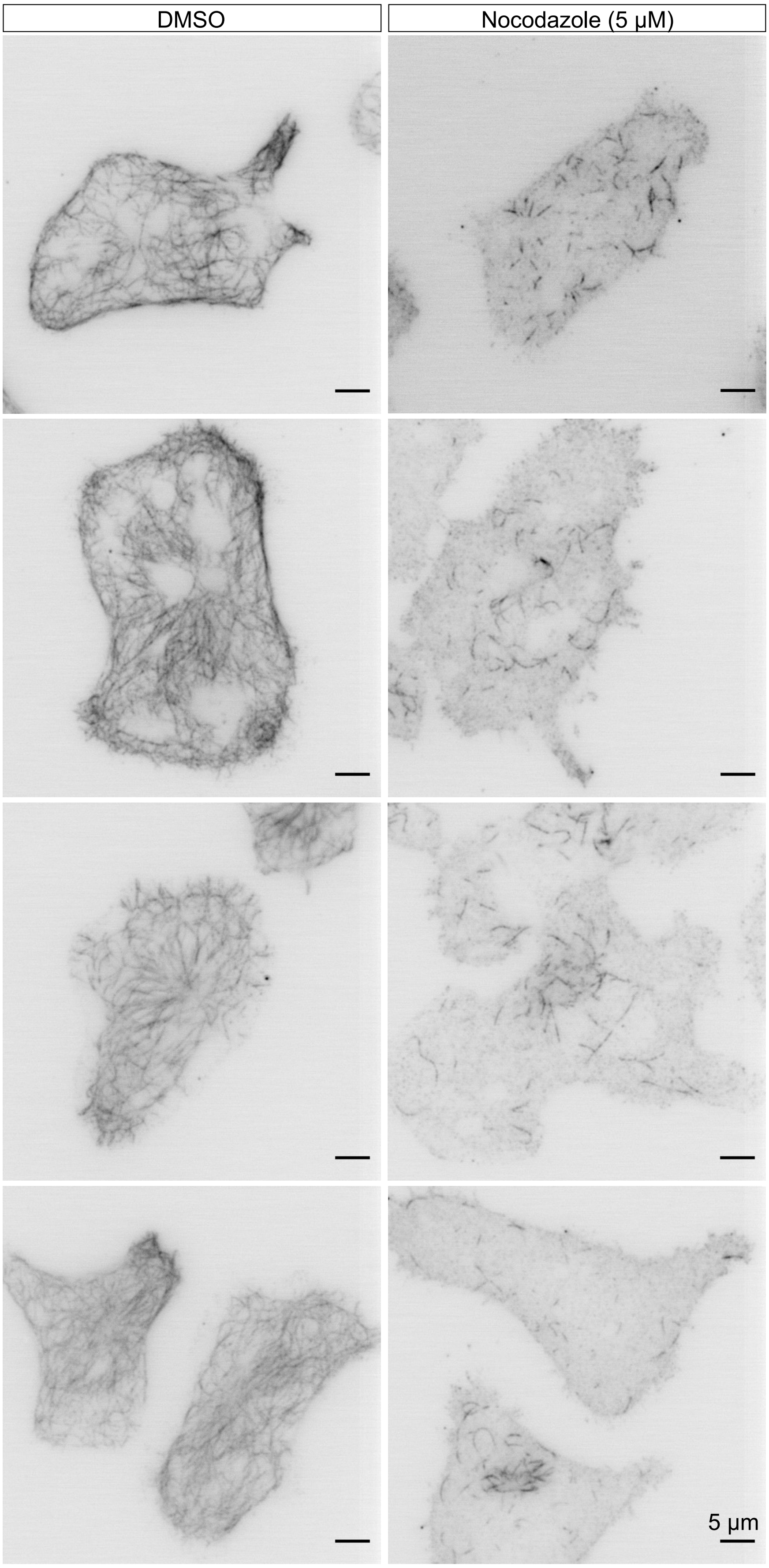


**Supplementary Figure 9. Nocodazole-treated cells have less microtubule filaments. a,** Low-resolution TIRF images of microtubules stained with anti-tubulin antibody in nocodazole-treated (left) and DMSO-treated (right) fixed cells acquired in the green channel.

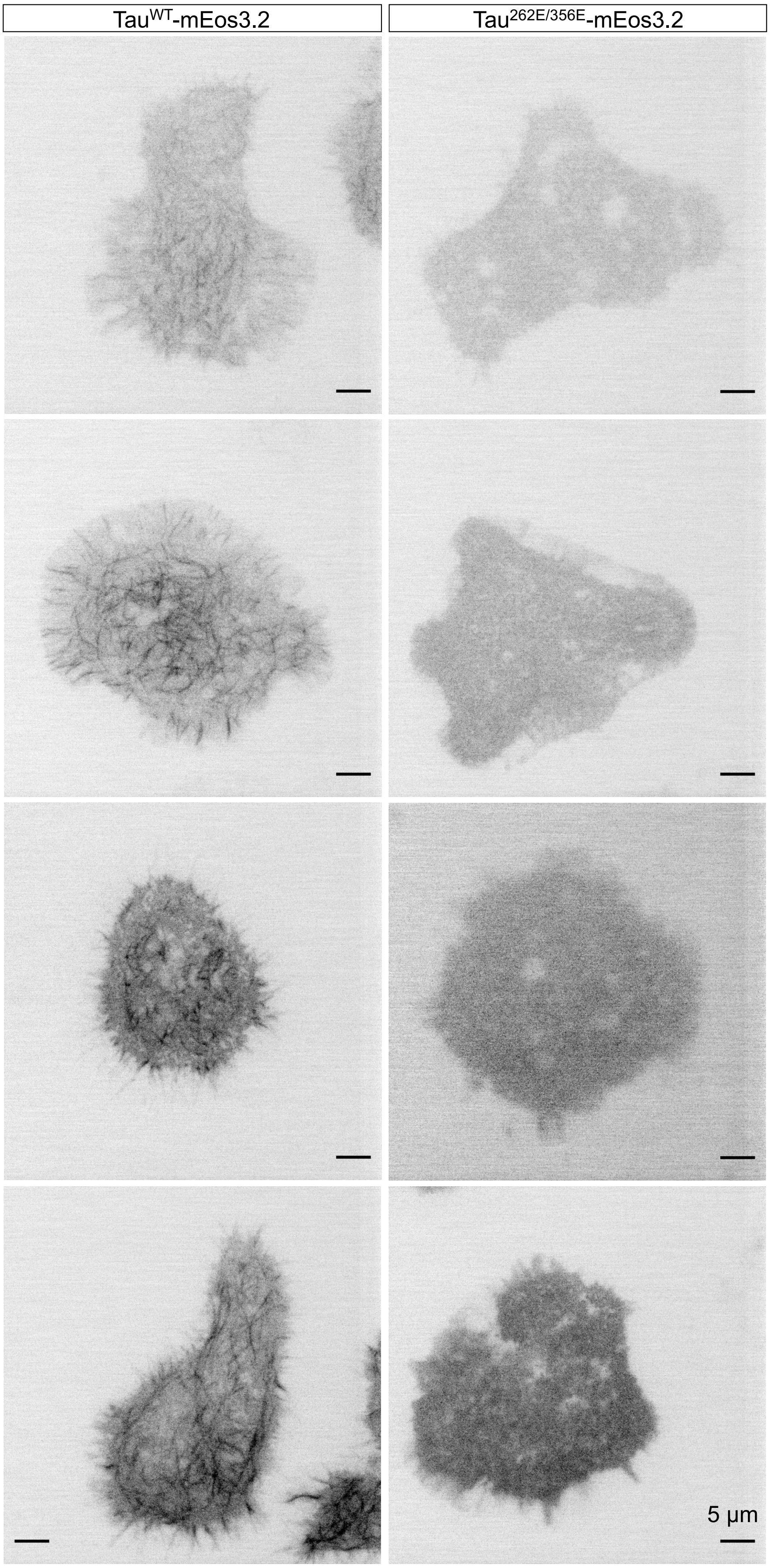


**Supplementary Figure 10. Microtubule-binding deficient mutant-expressing cells have less filament-like structures.** Low-resolution TIRF images of live cells expressing either $Tau^{WT}$-mEos3.2 (left) or $Tau^{S262E/356E}$-mEos3.2 (right) acquired in the green channel.

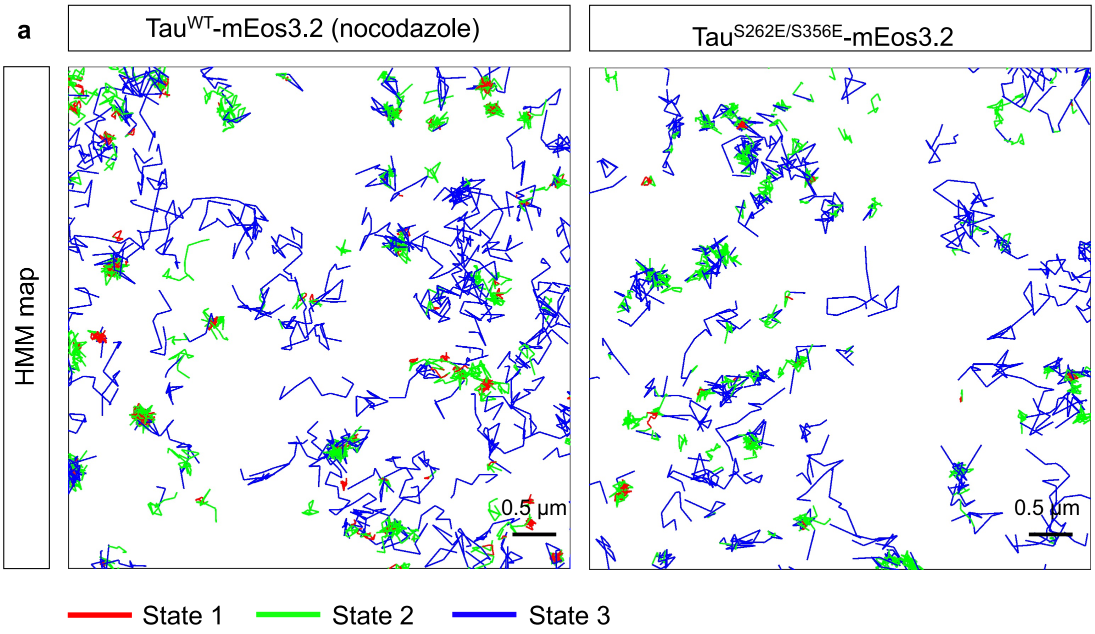


b

| | Tau$^{WT}$-mEos3.2 (nocodazole) (n=26 cells) | | | Tau$^{S262E/S356E}$-mEos3.2 (n=38 cells) | |
|---|---|---|---|---|---|
| | Apparent diffusion coefficient ($\mu m^2\ s^{-1}$) | State occupancy (%) | | Apparent diffusion coefficient ($\mu m^2\ s^{-1}$) | State occupancy (%) |
| State 1 | 0.043±0.009 | 27.9±5.4 | State 1 | 0.061±0.02 | 27.9±11.6 |
| State 2 | 0.116±0.031 | 41.1±4.0 | State 2 | 0.17±0.07 | 41.2±5.1 |
| State 3 | 0.31±0.02 | 31.0±6.3 | State 3 | 0.33±0.02 | 30.9±9.6 |

**Supplementary Figure 11. Tau displays multiple motion states even after preventing Tau/microtubule interactions. a,** Maps of Tau$^{WT}$-mEos3.2 trajectories in nocodazole-treated cells (left) and Tau$^{S262E/356E}$-mEos3.2 trajectories in untreated cells (right) annotated as distinct diffusive states using the 3-state hidden Markov model. The maps correspond to boxed regions highlighted in Figure 4. **b,** A summary of the apparent diffusion coefficients and the state occupancies estimated using the 3-state hidden Markov model. The mean±s.d. are shown.

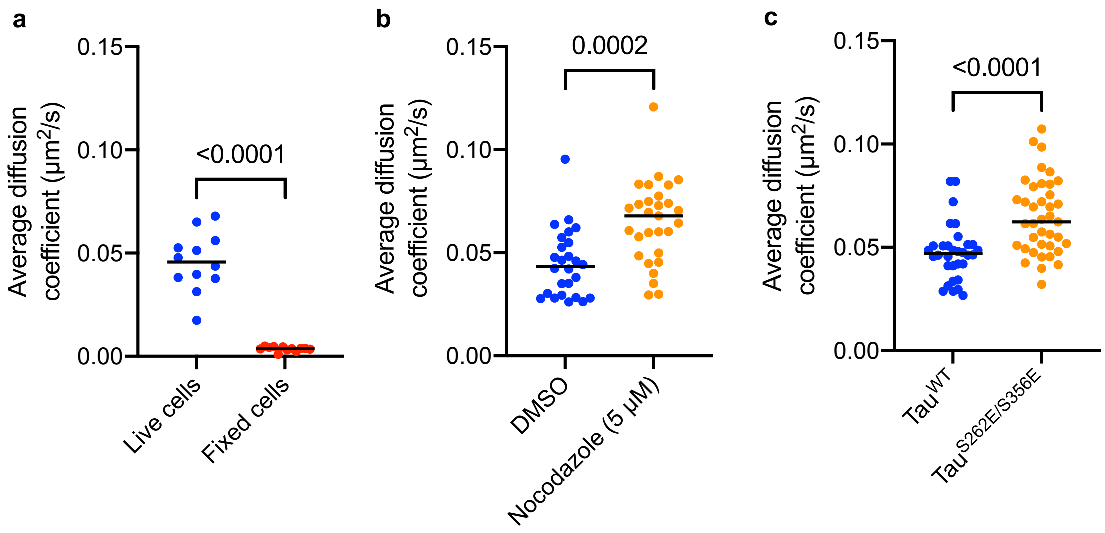


**Supplementary Figure 12. The average diffusion coefficient of Tau in different conditions. a-c,** The average diffusion coefficient of Tau corresponding to data in Fig. 1g (**a**), Fig. 4e (**b**), and Fig. 4n (**c**).
